# Equality between the sexes in plants for costs of reproduction; evidence from the dioecious Cape genus *Leucadendron* (Proteaceae)

**DOI:** 10.1101/212555

**Authors:** Jeremy J. Midgley, Adam G. West, Michael D. Cramer

## Abstract

It has been argued that sexual allocation is greater for female function than male function in plants in general and specifically for the large dioecious Cape genus *Leucadendron*. Here, we use new interpretations of published information to support the hypothesis of equality between sexes in this genus. The explanations are based on the fire ecology of the Cape that results in reproductive synchrony, reproductive doubling and competitive symmetry. Firstly, strict post-fire seedling establishment of the reseeder life-history in the Cape results in single-aged populations. Consequently, the reproductive and vegetative schedules of males must synchronously track that of females. This implies equal allocation to sex. Secondly, after fires, dioecious females have double the seedling to adult ratio of co-occurring hermaphrodites. This indicates that being liberated from male function allows females access to resources that double their fitness compared to hermaphrodites. Therefore, male and female costs of reproduction are equal in hermaphrodites. Thirdly, competitive symmetry must occur because males and female plants will frequently encounter each other as close near neighbours. Competitive asymmetry would both reduce mating opportunities and skew local sex ratios. The evidence to date is for 1:1 sex ratios in small plots and this indicates competitive symmetry and a lack of dimorphic niches. Finally, vegetatively dimorphic species must also allocate equally to sex, or else sexual asynchrony, lack of reproductive doubling or competitive asymmetry will occur.

## Introduction

Understanding allocation by the different sexes is a key focus in evolutionary studies and in organisms where both parents contribute genes equally to off-spring, sexual allocation (e.g. sex ratios, parental care of off-spring of different sexes, costs of reproduction) is expected to be equal (Fischerian) (Hamilton 1967; Charnov 1982). It is difficult to investigate sexual allocation in plants because most plants (>90%) are hermaphrodites, nevertheless, it is generally assumed that the resource costs of reproduction are greater for females (Barrett and Hough 2013). Although only female plants bear the costs of producing large seeds, fruits and cones after flowering, it is nevertheless controversial whether total female costs are higher than male costs. For example, Harris and Pannell (2010) argued that serotiny (storage of seeds in closed cones that open after plant death in fire) is a form of maternal care that imposes extra selection pressure on females of the Cape Proteaceae genus *Leucadendron* for greater water efficiency. They also argue that vegetative dimorphism, such as the larger leaves and lower rates of branching in females, results from this extra ecophysiological selection on females. Cones on females need to maintain a water supply to prevent them from drying out during the dry, hot Mediterranean summers and opening prematurely and thus releasing seeds into the unfavourable inter-fire environment. The proposed mechanism for greater water efficiency in females is the hydraulic benefit of their less frequent branching than occurs in males. Barrett et al. (2010) noted that males in serotinous *L. gandogeri* matured first and argued that the delayed maturation of females reflects their higher costs of reproduction.

In contrast to these suggestions of higher female costs, Midgley (2010) showed that δ ^13^ C (an integrated measure of an individual’s water use efficiency) did not differ between males and females of eight *Leucadendron* species, including a co-occurring vegetatively monomorphic and very dimorphic species, as well as serotinous and non-serotinous species. Midgley (2010) also argued that vegetative dimorphism in leaf size and branching is not explained by ecophysiological differences between the sexes, but differences in the process of pollen transfer and capture. Thus, whether the costs of sex are higher for females and whether this could lead to vegetative dimorphism is unresolved for *Leucadendron*. Here we use new interpretations of published data to further support the hypothesis that the costs of sex are equal amongst the sexes. Our hypothesis for sexual equality should apply generally to all plants and could be tested using some of the many hundreds of Cape dioecious species (such as > 300 Restionaceae species) and using dioecious species from other similarly fire-prone environments such as in Western Australia.

The hypothesis of higher allocation to reproduction in female plants, besides being of fundamental evolutionary interest, may also have a conservation implication for dioecious taxa such as *Leucadendron*. It has been argued that global change will lead to a decline in dioecious species (Hultine et al. 2016). A more stressed future environment (e.g. greater aridity) is argued to favour male survival if males allocate relatively more to vegetative growth and less to reproduction than females. Female biased mortality will then have negative population consequences such as male biased sex ratios.

The fire-prone vegetation of the Cape (*fynbos*), South Africa, is an excellent place to test the hypothesis of greater female allocation to reproduction; allocation to sexual reproduction rather than vegetative reproduction is critical for post-fire seedling recruitment, dioecy is common and the fitness of co-occurring dioecious and hermaphroditic taxa can be compared. Also, sexual dimorphism in the genus *Leucadendron* (Proteaceae) can be extreme at a global scale.

### The fynbos environment

*Leucadendron* contains about 80 species and occurs in fire-prone Cape (South Africa) vegetation known as *fynbos* (*sensu* Rebelo et al. 2006). The Proteaceae dominate the fynbos canopy layer which is about 2 m tall, mainly through the genera *Protea* and *Leucadendron*. The family (about 360 fynbos species) includes both dioecious (25%) and hermaphroditic taxa and which frequently co-exist Figure 1). Most Proteaceae species are reseeders (i.e. non-resprouters) (Le Maitre and Midgley 1992). Since reseeders cannot resprout, all plants die in fires. As they are relatively short-lived (1-3m tall, 5-30 years), plants also die in senescent vegetation which occurs in the prolonged absence of fire (Bond 1980). Successful recruitment of reseeders only takes place after a fire (Bond et al. 1987; Le Maitre and Midgley 1992). About 60% of fynbos Proteaceae species are serotinous (Le Maitre and Midgley 1990). When serotinous cones dry-out and open after death of adults in fire, seeds are released en masse into the optimum post-fire environment where seedling recruitment takes place in situ Figure 2). The most common soil seed-bank (35% of fynbos Proteaceae species) is due to ant burial of seeds (Slingsby and Bond 1985; Le Maitre and Midgley 1992) and most of remaining 5% of species have seeds that are scatter-hoarded by small mammals (Midgley and Anderson 2005). Species with soil seed-banks also germinate after fire as do serotinous species; except in this case typically fire is needed to crack their relatively thick seed coats. All three of these types of seed-stores occur in *Leucadendron*.

**Figure 1.**
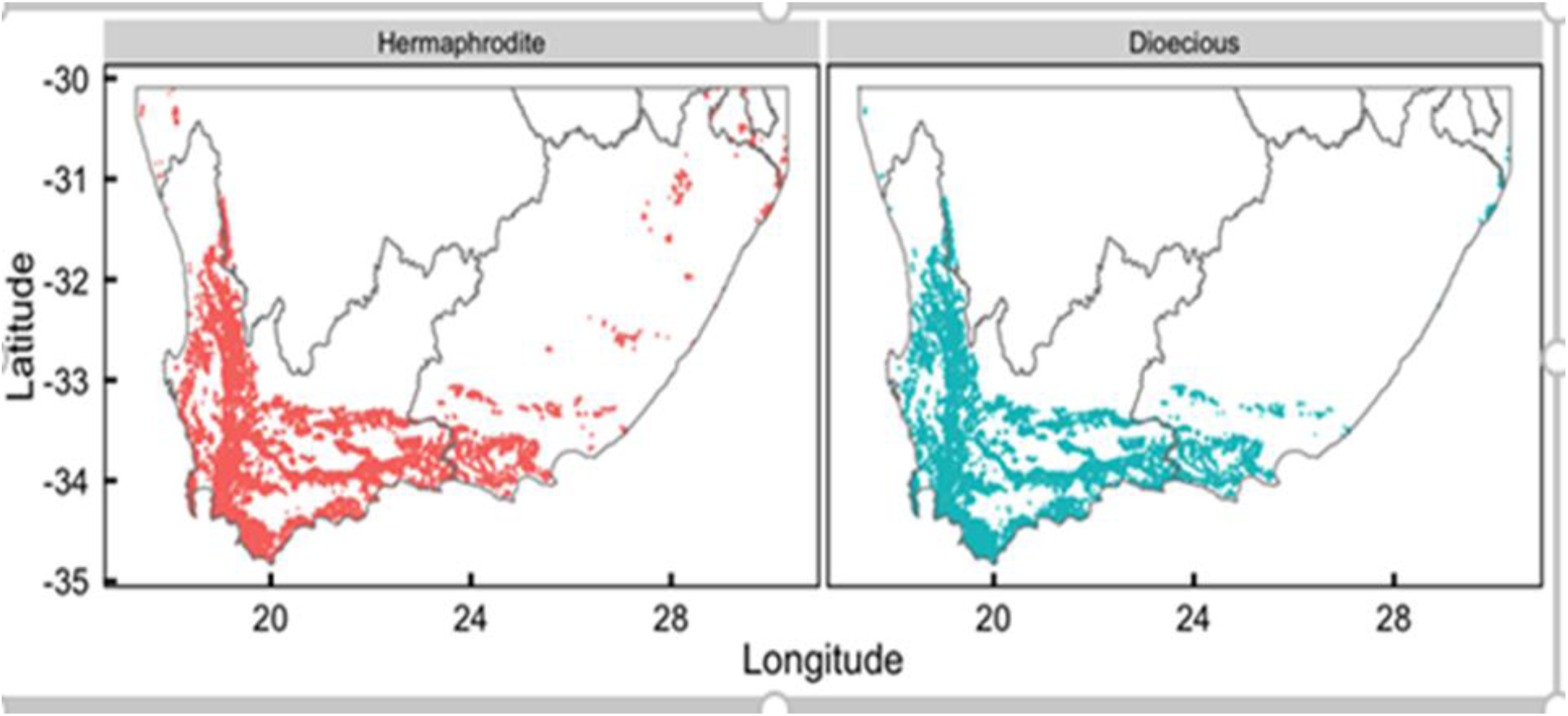
Distribution of hermaphrodite (*Brabejum, Diastella, Faurea, Leucospermum, Mimetes, Orothamnus, Paranomus, Protea, Serruria, Sorocephalus, Spatalla* and *Vexatorella*) and dioecious genera (*Leucadendron* and *Aulax*) obtained from the location records for Proteaceae in Protea Atlas dataset (www.proteaatlas.org.za). The coastline and provincial borders are shown for reference.

**Figure 2.**
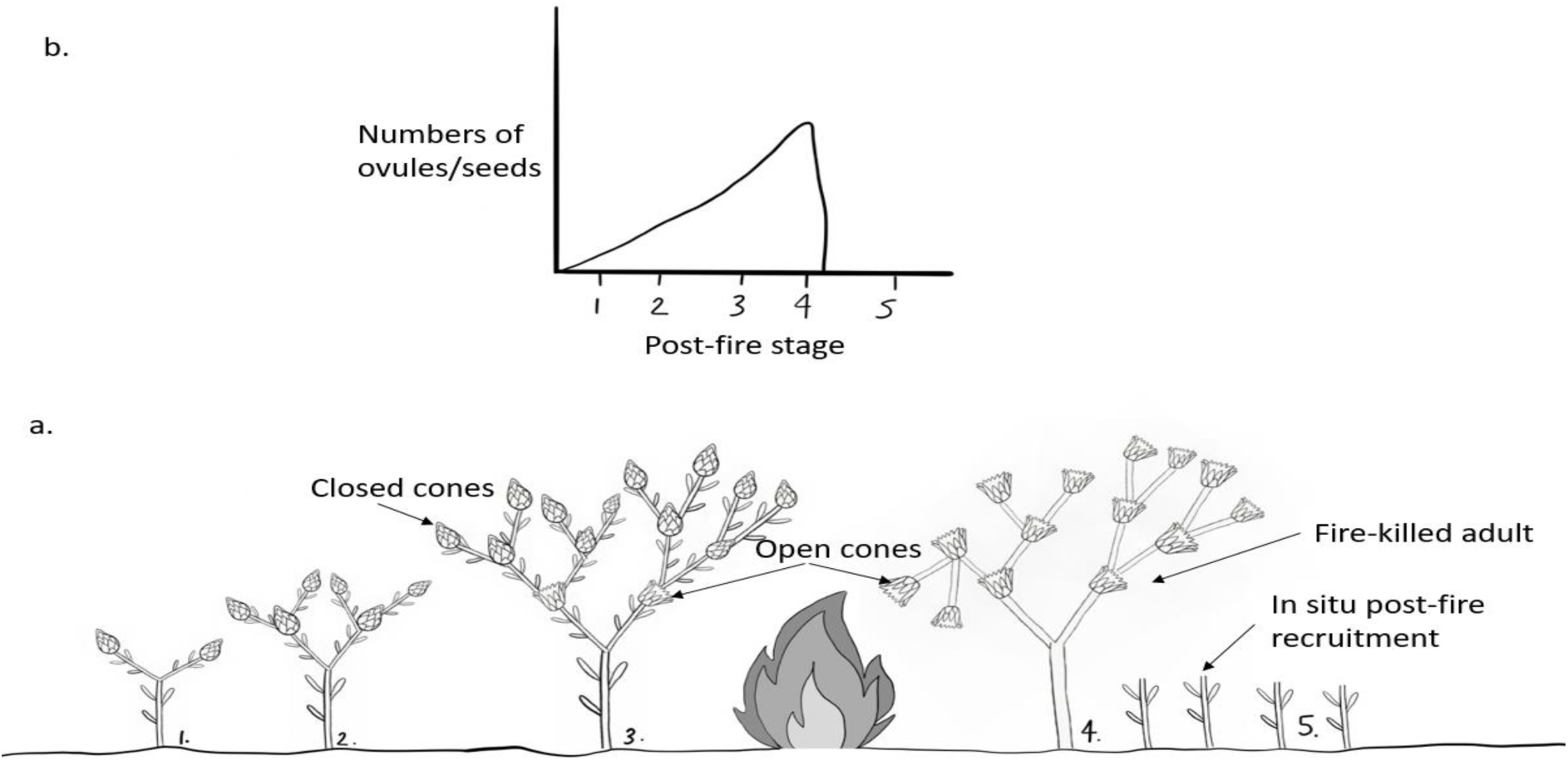
a. Stylised life-history of a serotinous reseeding proteoid. Phases 1-3 depict rapid increase of seeds over post-fire time. Cones remain closed for about two years and most cones open simultaneously after fire (Phase 4) and release seeds. Single-aged cohorts of seedlings establish in situ post-fire (Phase 5). b. Rapid increase in number of ovules over post-fire phases for males to compete for, and numbers of seeds available for post-fire recruitment.

Given that only fires create suitable opportunities for regeneration, there is no strong selection in fynbos for long distance seed dispersal such as by birds (Le Maitre and Midgley, 1992). Plant seeds merely wait in situ in soil or canopy seed-banks for recruitment opportunities post-fire. Dispersing long distances could result in hazardous movement away from recently burned sites to unsuitable unburned sites.

Short distance seed dispersal thus predominates in the fynbos Proteaceae and is due to both the dispersal agents and short plant size. Scatterhoarding by small mammals (Midgley and Anderson 2005) and ant-dispersal (Slingsby and Bond 1985) results in dispersal at the scale of metres. Dispersal of winged serotinous *Leucadendron* seeds has not been studied but is also expected to be very local, given that they are released from very short plants (0.5-3 m tall) and that release height dominates wind dispersal distances (Augsberger et al. 2016). Wind dispersed *Leucadendron* species have small wings (Williams, 1972) and relatively high fall-rates (Midgley et al. in prep.). Genetic studies indicate median dispersal distance in relatively large winged *Banksia* (Proteaceae) seeds is only 5 m (Krauss et al. 2009). Some species from both hermaphroditic and dioecious taxa produce seeds that are hairy and can be wind-rolled along the soil but are rapidly captured by obstacles and depressions (Bond 1988, 95 % of seedlings < 15 m from parent plant in Manders 1986). The features of the Proteaceae in the fynbos environment that are relevant to considering sexual costs of reproduction are; (i) co-occurring dioecious and hermaphrodite species, (ii) single-aged populations of the reseeder life history and (iii) short gene (see pollination aspect later) dispersal distances, which render post-fire recruitment patterns as a measure of local dead adult fitness. Below we argue why these characteristics should result in equal reproductive allocation between the sexes of dioecious plants.

### Reproductive synchrony; *Leucadendron* populations are single-aged and this forces males to track females

Fynbos reseeder populations are single-aged, dating back to winter germination after the most recent fire. Individual female reproductive schedules will evolve in concert with the historical fire regimes such that maximum output coincides with the most likely fire frequency. The historical variance around the mean fire frequencies would constrain age to maturity and senescence, because reseeders need to be reproductive if fire occurs in young vegetation and serotinous females need to be alive in old vegetation. Since fynbos Proteaceae grow and mature within a few years (Midgley and Rebelo 2010) and because serotiny is generally weak (cones are typically only retained for a year or two, seed viability declines with time), the cumulative seed crop is dominated by the contribution of the most recent few years (Midgley, 2000). Therefore, given no immigration by long distance dispersal, the level of post-fire seedling recruitment is largely dominated by the reproductive condition of the females at the time of the fire, Figure 2). For example, fire in very young or senescent vegetation results in very low dead adult:seedling ratios (e.g. Bond 1980; Kraaij et al. 2013).

The scheduling of male reproductive allocation in single-aged reseeders must synchronously match that of females, whatever the factors are that constrain the scheduling female reproductive allocation. Males that allocate more to sexual function and less to vegetative function at an earlier age than females would experience limited mating opportunities due to being asynchronous with ovule numbers for their whole lifetime. Similarly, a male that allocated more to vegetative growth than females to achieve larger size at maturity would also be asynchronous with females. Also, large relatively immature males forego future breeding opportunities when a fire takes place because fires kill all individuals. Synchronous allocation to reproduction also implies equal allocation to vegetative growth as there are no advantages of delayed maturity to achieve relatively larger size or longevity, given that fire kills all plants.

### Reproductive doubling; post-fire recruitment levels of dioecious females is double that of hermaphrodites

Because adults die in fires and seed dispersal distances are short, local extinction will result after fire in a stand of immature or senescent plants. Post-fire recruitment levels (i.e. dead adults:seedling ratios) of the Proteaceae have therefore been widely measured in fynbos, to determine responses of the reseeders to fires in stands of different age and in different seasons (Bond 1984; Bond et al. 1984; Midgley 1987; Kraaij et al. 2013). These surveys are typically determined 1-4 years after fires. However, since most fynbos Proteaceae species grow and mature rapidly (generally within 1-5 years, Midgley and Rebelo 2008, Kraaij et al. 2013) these recruits are not seedlings (i.e. dependent on seed reserves) but can be advanced juveniles that have already survived summer drought and competitive interactions. Given the predominance of short distance dispersal, we argue that post-fire dead adult:seedling ratios of reseeders are therefore a direct measure of mean lifetime fitness and that this then allows fitness comparisons between co-occurring dioecious and hermaphroditic taxa. Since dioecious and hermaphrodites co-occur, mean fitness must be similar.

Species from fynbos dioecious genera frequently co-occur in the same landscape, even the same plots with those from similar sized species from hermaphroditic genera; *Leucadendron* and *Aulax* co-occur with hermaphroditic genera *Protea*, *Leucospermum* and *Mimetes* (Supplementary Figure 1 and see also community descriptions in Rebelo et al. 2006). Because dioecy is rare amongst plant species (< 5%), it is often considered to be disadvantageous, although this is questioned by Sabath et al. (2016). In fynbos, the widespread co-occurrence of dioecious and hermaphroditic taxa indicates there is no nett benefit, or disadvantage, of either breeding system. Thus, proteas and leucadendrons are ecological equivalents given their fine-scale co-occurrence as well as other forms of equivalence such as in plant size, reseeding life-history, serotiny and seedling growth rates (Laurie et al. 1997). This ecological equivalence and stable co-occurrence means that they must both have the same mean post-fire fitness (i.e. dead adult:seedling ratio). For example, if the female dioecious female adult:seedling ratio is double the hermaphrodite adult:seedling ratio, this implies a 1:1 sex ratio in the dioecious taxa.

Post-fire recruitment data indicates dioecious female individuals do produce double the number of seedlings as do co-occurring hermaphrodites (Midgley 1987), such that the standard measurement of dioecious recruitment levels is female adults*2/seedlings (e.g. Kraaij et al. 2013). For a recent example, our analysis of the post-fire recruitment data in Kraaij et al. (Fig. 4, 2013) indicates there is no difference between in seedling:adult ratios in *Protea* versus seedling:2*female ratios in *Leucadendron* (M-W p >0.2). Dioecious females achieve higher seedling recruitment in part by having higher seed set and thus seed production than hermaphrodites. For example, seed set (from Mustart et al. 1994) in co-occurring *Protea obtusifolia* (19.5%) is lower than in female *L. meridianum* (54.7%) as is *P. susannae* (12.7%) compared to co-occurring *L. coniferum* (87.1%).

The doubling of female seedling production in *Leucadendron* is not because low seed production in hermaphrodites is limited by out-crossed pollen. Most of the hermaphroditic Proteaceae are also out-crossers (Collins and Rebelo 1987), mainly through strong protandry and self-incompatibility. Despite the out-bred genetic structure of most hermaphroditic Proteaceae (e.g. Goldingay and Carthew 1998), the widely observed low seed-set (<25%) is not strongly improved by artificially adding cross pollen (e.g. Whelan and Denham 2009; Steenhuisen and Johnson 2012). This generally low seed in the Proteaceae is not maladaptive, it reflects functional andromonoecy (Rebelo and Collins 1987); most flowers/florets have evolved to achieve fitness by male function and the distribution of pollen. Support for the functional andromonoecy hypothesis is the higher seed set (> 50%) in female florets in dioecious taxa such as in *Aulax* and *Leucadendron* (Rebelo and Collins 1987; Mustart et al.1997; Hemborg and Bond 2005), but that this is normalised when lack of seed-set in the many male florets on male plants is considered. Also, seed-set is very high in the hermaphrodite flowers in morphologically andromonoecious reseeding proteoids, but it too is very low when lack of seed-set in the numerous male flowers per plant is considered (Ladd and Connell, 1994). In the few somewhat self-compatible Proteaceae species that do occur, seed-set per floret is still very low (< 1-3 % in several stands of *Banksia baxteri*) (Wooller and Wooller 2001). In this species, seed-set and seed quality is much higher in open controls than manually-selfed flowers (Wooller and Wooller 2004), indicating that most seed-set in open flowers is due to out-crossed pollen. Furthermore, Hemborg and Bond (2005) noted that 95% of female *L. xanthoconus* flowers were pollinated, but only about 50% developed into seeds. In summary, nett seed-set in both dioecious and hermaphroditic taxa is similar and in neither, is specifically out-cross pollen limited.

Double *Leucadendron* female fitness (dead adult:seedlings) compared to co-occurring hermaphrodites indicates that male and female costs of reproduction in hermaphrodites are equal; a dioecious female liberated from male function can produce twice as many recruits as a hermaphrodite. Double female fitness also indicates that the adult sex ratio in dioecious plants is 1:1; a dioecious male and female have the same fitness as two hermaphrodites. The above reproductive argument applies to male components of fitness, but it is more difficult to measure than is the seedling ratios of females. For example, burned males do not have cones which thus prevents them being identified and counted.

Our argument for equal costs of male and female function appears to run counter to arguments of differential resource utilisation by the sexes. For example, it has been argued that males and females use different resources for reproduction, such as nitrogen for pollen and carbon for seeds (Harris and Pannell, 2008). However, the doubling of seedling production of dioecious females relative to that of hermaphrodites indicates that the release from the cost of male function must be equal in magnitude to that of the female function.

Thus, even if males and females use resources at different stoichiometries, the nett cost of these functions to the plant must be equal.

### Competitive symmetry is needed to maintain 1:1 sex ratios

Short distance seed dispersal from large seed-banks will cause intense post-fire competitive interactions. Numbers of seeds stored in individual serotinous adult *Leucadendron* canopies can be large (e.g. > 24 000 per female in *L. coniferum* and > 2500 in *L. meridianum* in Mustart et al. 1994). Since fynbos Proteaceae often occur in mesic areas, dense stands of seedlings typically arise after fire in mature stands. The number of 1-4-year-old seedlings within the space occupied by an adult is up to 50 in Bond et al. (1987) and 40 in Kraaij et al. (2013), with densities up to > 200 m^−2^ in *L. uliginosum* (Midgley 1987). In many animal species, competition between sexes can be avoided, but not in dense stands of plants. Thus, many potential competitive intra-specific near neighbour interactions will occur in *Leucadendron*. In dioecious fynbos Proteaceae, seed dispersal attributes do not differ between the sexes because the dispersal attributes (elaiosome, wings or hairs) are maternal (i.e. they are fruits, Midgley 1987), not off-spring tissue. Given equal dispersal of an equal ratio of male and female seeds, 50% of seedling interactions would randomly be mixed-sex encounters and 50% same-sex encounters. If males allocated less to reproduction than females, they would be vegetatively superior and out-compete females, thus causing local non 1:1 sex ratios and local mate competition.

Many entomophilous *Leucadendron* species are pollinated by very small (< 2mm in length, Williams, 1972) largely sedentary nitidulid insects (Hemborg and Bond 2005; Welsford et al. 2014). Thus, biased sex ratios may complicate pollination by increasing between sex distances that very small insects must travel. A male that allocates less to sexual reproduction and more to vegetative growth than conspecifics could outcompete near neighbour males and females. However, for this large male mutant that is capable of out-competing near neighbour females to be successful, it still faces the hurdle of dispersing pollen to other more distant females which will likely have smaller, but closer, near-neighbour males. This is unlikely because to be vegetatively competitive this mutant male must be allocating less to reproduction. Also, the spread of such an aggressive male genotype would in any event eventually be terminated by the strong decline in females (leading to 1:3 sex ratio) and the necessary evolution of compensatory female biased sex ratios of seeds (> 2:1), in contradiction of the evolutionary stable 1:1 sex ratio strategy for most dioecious organisms and the ratio expected and found in *Leucadendron*. Barrett et al. (2010) documented 1:1 sex ratios in mature mesic *Leucadendron* stands and analysis of the data of Mustart et al. (1994) indicates no difference in sex ratio from 1:1. Given the argument above that competitive interactions between the sexes are highly likely in fynbos, this sex ratio equality implies competitive symmetry across the sexes. This implies resource use is equal between the sexes.

### Serotiny, higher female costs and vegetative dimorphism

Besides the above arguments for equality between the sexes, there are theoretical and mechanistic reasons why the argument of Harris and Pannell (2010) based on extra maternal costs of serotiny cannot explain leaf dimorphism. Firstly, maternal care is not an extra cost compared to male sexual allocation, but a cost that must be integrated into female reproductive allocation. Storing seeds in canopies is merely an alternative adaptation to soil-stored seeds as a mechanism for recruiting after fynbos fires. For example, many scatterhoarded *Leucadendron* species have large seeds with a thicker seed coat, than is the case with serotinous species (Williams 1972; Midgley et al. 2015). The thick seed coat of these species is an alternative allocation to a cone, and cone maintenance, not an extra cost.

Cones remain closed for many years in the serotinous dioecious Proteaceae genus *Aulax*, but according to the leaf size data in Rourke (1987), there is no sexual leaf dimorphism. Intense serotiny is therefore not a general correlate of dimorphism. Leaf dimorphism in *Leucadendron* matches floral dimorphism via Corner’s Rules (Bond and Midgley 1988); few, large inflorescences on fewer larger branches in females versus many, smaller inflorescences and branches in males. Explaining dimorphism based on water relations cannot explain why then males have gone the apparently expensive “many-small” route rather than also evolving this more effective habit (Midgley 2010). Midgley (2010) argued that the “many-small versus few-big” is due to differences in dispersing and receiving pollen.

Regarding ecophysiological mechanisms for the evolution of dimorphism, the hydraulic advantages of larger leaf size and lower branching are controversial. Despite the allometric correlations summarised in Corner’s Rules being important in plants (e.g. Westoby et al., 2002), there has been no argument that lower rates of branching have a positive hydraulic effect. Lower branching associated with larger leaf size *per se* is not predicted to make a substantial positive impact on carbon and water economy, because larger leaf size has other balancing positive and negative correlates. Larger leaf (and associated branch) size is correlated with lower specific leaf area (e.g. Midgley 2010 for *Leucadendron*), more self-shading and a longer path-length, but a larger xylem diameter and smaller total leaf area (West et al. 1999; Westoby et al. 2002; Smith et al. 2017). Within *Leucadendron* mean leaf size is smaller in the more arid areas (Thuiller et al. 2004), in contradiction of the expectation of Harris and Pannell (2010). Although cones transpire and respire at a total rate equivalent to only a few leaves (Cramer and Midgley 2000), transpiring cones must nevertheless compete with transpiring leaves on the same plant for water (Midgley and Enright 2000). Even if reduced branching rates was positively correlated with hydraulic efficiency, it would equally influence all leaves and cones on the same plant and thus not improve relative access to water of cones of different ages. If lower branching patterns positively influenced plant water status, then this should also apply to non-serotinous species as well. Finally, Bond and Midgley (1988) noted leaves on *Leucadendron* seedlings resemble those of females, suggesting that leaf dimorphism is primarily due to selection on males, not females. In conclusion, there is no expectation to the contention that lower branching/larger leaves is positively associated with water use efficiency and measurements showed no sexual differences in water use efficiency (Midgley 2010).

### Male costs of reproduction in *Leucadendron*

The large investment in cones and seeds by females tends to dominate the view of higher female’s costs of reproduction. However, if male and female reproductive costs are the same, where do the high male’s costs come from? There are features of male reproduction in *Leucadendron*, besides the cost of pollen, that incur higher costs than in females. During flowering in *Leucadendron* the bracts and leaves surrounding male inflorescences of insect pollinated species are far more yellow and less green than females; in many species, the whole male plant may become yellow/red, presumably to attract pollinators. This sexual dimorphism in attractiveness is due to a total break-down of thylakoids in leaves and bracts, which then exposes hidden yellow carotenoids (Schmeisser et al. 2010). This break-down of chlorophyll and the accompanying loss of photosynthesis, must be far more expensive to males than females because it occurs in many more leaves and floral bracts in male plants Figure 3). Males produce more scent and with a greater diversity of volatiles than females (Welsford et al. 2015). Also, males in wind-pollinated *Leucadendron* species produce many more florets than do their females (Midgley 2010). Thus, male allocation to reproduction is as high as in females; through pollen, greater attractiveness and high male:female floret ratios.

**Figure 3.**
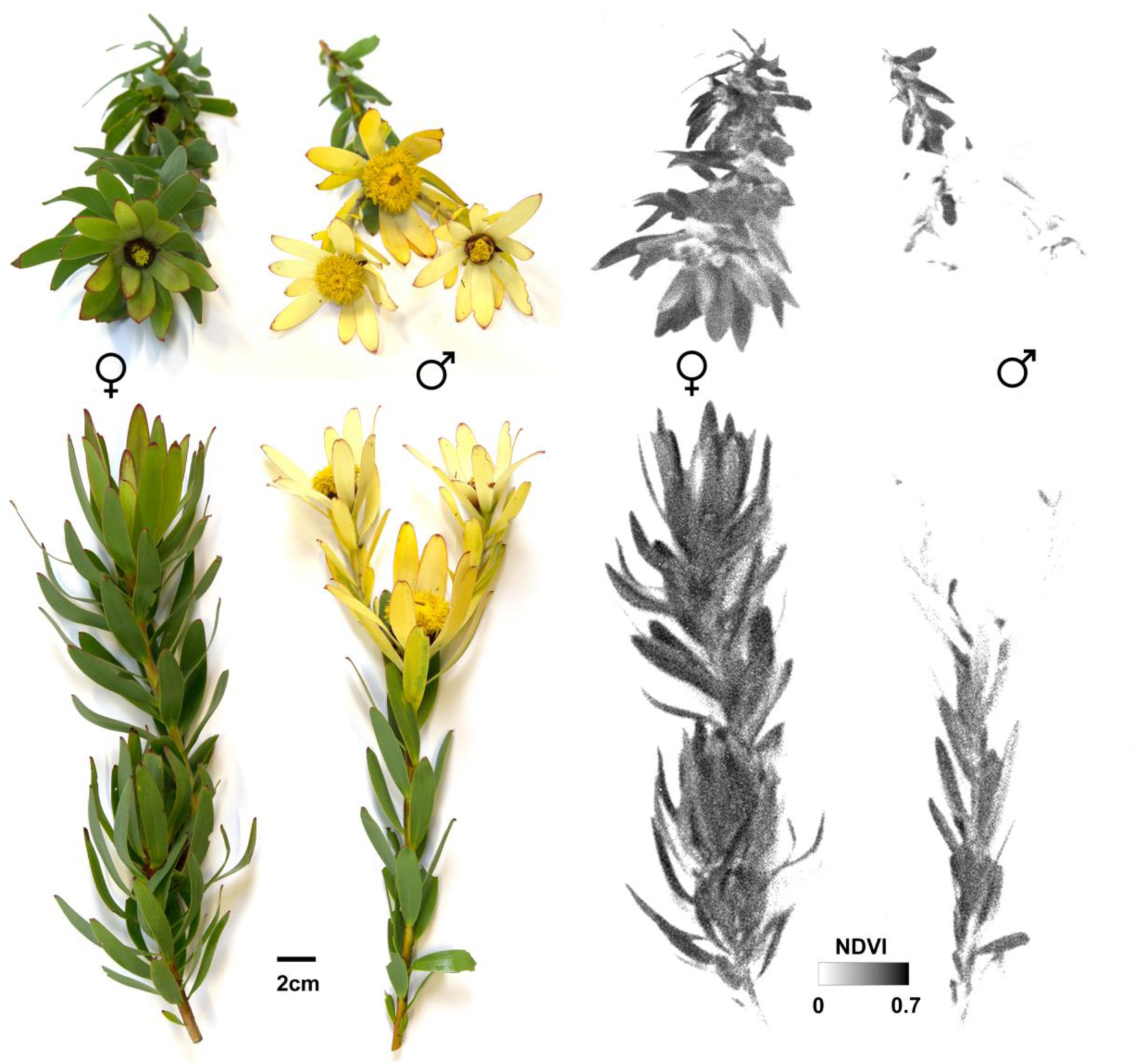
Representative male and female flowering shoots of *Leucadendron sessile* in colour (left) and Normalised Difference Vegetation Index (right). NDVI image created using a Parrot Sequoia multispectral camera. Note low NDVI in more male terminal leaves and bracts than females.

### Sexual niche segregation is unlikely

Females could sustain higher reproductive allocation if they out-competed males in more resource rich sites. This is often seen in riparian ecosystems where steep and stable environmental gradients occur over short spatial scales, allowing sexual niche-segregation and specialisation (e.g. Hultine et al. 2008). However, this is unlikely to be the case in the fynbos for several reasons. We have already argued that in fynbos dioecious males will always track female allocation, so this competitive elimination of one sex by the other in some habitats is unlikely. Furthermore, we suggest that frequent fire, short lifespans and limited gene dispersal in space and time by either seeds or pollen reduces the opportunities for the sexes to specialise in different microsites. Firstly, such microsites would have to be at a very fine geographic scale (a few metres) in the fynbos to match the limited gene dispersal patterns. Secondly, frequent fire resets sexual spatial patterns in this landscape because all adults die in fire, but seed dispersal of both sexes only occurs from female plants. For a male mutant genotype that is advantageous in an arid microsite to succeed, this genotype must repeatedly disperse in seeds away from maternal plants in the mesic sites, out-compete females in the arid sites and then disperse in pollen back from the arid sites to females in the mesic sites, whilst competing with the mesic adapted males for mating opportunities. Furthermore, these sites must be burned in the same fires or else populations of each sex will be at different stages of reproductive maturity. This back-and-forward scenario from females after every fire is very unlikely. Finally, the data available suggest no habitat specialisation; males and females co-occur equally and at a very fine scale (such as in 5 × 5 m plots in Barrett et al. 2010 and 10 × 10 m plots in Mustart et al. 1994).

Even if there were ecophysiological consequences of vegetative dimorphism, say of differences in leaf size or hydraulic architecture, this would not change how present dimorphic females experience the past benefit of avoiding allocation to male function. This would have been set when dioecy first evolved and per the above arguments appears to have been a 50% reproductive benefit. Therefore, dimorphism in *Leucadendron* cannot be due to differences in sexual allocation because reproductive asynchrony, lack of reproductive doubling or competitive asymmetry would then occur. We have argued above that sexual niche dimorphism is unlikely and therefore sexual vegetative dimorphism cannot be a result of sexual niche dimorphism. Vegetative dimorphism is therefore not due to ecophysiological differences, habitat differences nor to allocation differences but could be due to differences in sexual efficiency related to pollination (Midgley 2010).

### Early maturation of males does not necessarily reflect higher costs of reproduction

We interpret the earlier maturation of *Leucadendron* males (Barret et al., 2010) not to represent sexual differences in total reproductive allocation but to differences in costs of producing individual inflorescences. Given that Cape Proteaceae are single-aged there would be strong selection preventing males from flowering if no females were flowering. Male inflorescences are far smaller than females (e.g. 0.6 % female mass in *L. rubrum* and 2.5% in *L. ericifolium and L. teretifolium* from Bond and Midgley 1988). Given that only one inflorescence needs to be produced to determine the sex of a plant and that size thresholds for maturity are widespread in plants, it is to be expected that the sex producing the cheaper *single* inflorescence will mature first. Although about double the number of males than females have inflorescences in 6-year-old stands of *L. gandogeri* (Barret et al. 2010), because females nevertheless have many more florets per single inflorescence than males, precocious male flowering is not maladaptive nor evidence of lower male total allocation to reproduction.

## Conclusion

Three separate arguments support the hypothesis that the costs of reproduction are equal in *Leucadendron*. Single-agedness forces males to allocate to reproduction in synchrony with females. Dioecious females achieve double the female fitness of hermaphrodites by being relieved of male function. Competitive interactions are likely in fynbos and since the data suggests 1:1 sexual ratios this indicates competitive symmetry. We conclude also that the costs of reproduction must be equal whether the taxa are dimorphic or not. Our argument applies in the Cape, where allocation to sexual reproduction is critical and where dimorphism can be extreme, but should be tested globally.

We predict that global change will impact dioecious Proteaceae to the same extent as hermaphrodites, because the latter being bi-parental and largely out-crossed, also allocate equally to the sexes. Plants from the different sexes in dimorphic species in *Leucadendron* have very different shapes and sizes. We nevertheless predict that whole plant resource use, sex ratios and near neighbour associations will be equivalent between the sexes within both monomorphic and dimorphic species and will not be differentially impacted by global change.

## Authors Contribution

JJM, AGM and MDC were all involved in all stages of this project.

## Acknowledgements

We thank Bruce Anderson, William Bond, Richard Cowling, Todd Dawson, Allan Ellis, Steve Johnson, Phil Ladd, Peter Linder, Laurence Midgley, John Pannell, Tony Verboom and Joe White for critical opinions.

